# Analysis of novel domain-specific mutations in the zebrafish *ndr2/cyclops* gene generated using CRISPR-Cas9 RNPs

**DOI:** 10.1101/277715

**Authors:** Ashley N. Turner, Reagan S. Andersen, Ivy E. Bookout, Lauren N. Brashear, James C. Davis, David M. Gahan, John P. Gotham, Baraa A. Hijaz, Ashish S. Kaushik, Jordan B McGill, Victoria L. Miller, Zachariah P. Moseley, Cerissa L. Nowell, Riddhi K. Patel, Mia C. Rodgers, Yazen A. Shihab, Austin P. Walker, Sarah R. Glover, Samantha D. Foster, Anil K. Challa

## Abstract

Nodal-related protein (ndr2) is a member of the transforming growth factor type β superfamily of factors and is required for ventral midline patterning of the embryonic central nervous system in zebrafish. In humans, mutations in the gene encoding nodal cause holoprosencephaly and heterotaxy. Mutations in the *ndr2* gene in the zebrafish (*Danio rerio*) lead to similar phenotypes, including loss of the medial floor plate, severe deficits in ventral forebrain development, and cyclopia. Alleles of the *ndr2* gene have been useful in studying patterning of ventral structures of the central nervous system. Fifteen different *ndr2* alleles have been reported in zebrafish, of which eight were generated using chemical mutagenesis, four were radiation-induced, and the remaining alleles were obtained via random insertion, gene targeting (TALEN), or unknown methods. Therefore, most mutation sites were random and could not be predicted *a priori*. Using the CRISPR-Cas9 system from *Streptococcus pyogenes*, we targeted distinct regions in all three exons of zebrafish *ndr2* and observed cyclopia in the injected (G_0_) embryos. We show that the use of sgRNA-Cas9 ribonucleoprotein (RNP) complexes can cause penetrant cyclopic phenotypes in injected (G_0_) embryos. Targeted PCR amplicon analysis using Sanger sequencing showed that most of the alleles had small indels resulting in frameshifts. The sequence information correlates with the loss of ndr2 activity. In this study, we validate multiple CRISPR targets using an *in vitro* nuclease assay and *in vivo* analysis using embryos. We describe one specific mutant allele resulting in loss of conserved terminal cysteine-coding sequences. This study is another demonstration of the utility of the CRISPR-Cas9 system in generating domain-specific mutations and provides further insights into the structure-function of the *ndr2* gene.

## Introduction

The transforming growth factor β (TGF-β) superfamily is one of the major groups of secreted signaling molecules that is important in cell-to-cell communication and coordinating pattern formation during development [1, 2]. The Nodal proteins of the TGF-β superfamily have been found in every vertebrate examined to date. Nodal factors play pivotal roles during embryogenesis of chordates and have been implicated in several developmental processes, including mesoderm and endoderm formation, anterior-posterior patterning and left-right axis formation [3-8].

In humans, there are several documented mutations in the *NODAL* gene that cause holoprosencephaly and heterotaxy [9]. In zebrafish, there are three *NODAL-* related paralogs (*ndr1*/*squint, ndr2*/*cyclops*, and *ndr3/southpaw*) that are very similar but have specialized functions. During embryogenesis *cyclops* (*cyc*)*/nodal related 2* (*ndr2*) is required for ventral midline patterning of the central nervous system [3, 4, 7, 8, 10, 11] (henceforth referred only as *ndr2*). Mutations in *ndr2* in the zebrafish lead to similar phenotypes observed in humans, including loss of medial floor plate, severe deficits in ventral forebrain development, and cyclopia [3, 4, 10, 12-18].

The TGF-β proteins, including nodal proteins, are synthesized as precursors and the propeptide is cleaved off by proprotein convertase at a unique cleavage site [19]. These cleavage site sequences vary among TGF-β precursors [20]. One conserved feature consists of a dibasic RXXR sequence [1, 19, 21], while other recognition sequences that contain an additional basic residue (RXRXXR, or RXK/RR) are more rapidly cleaved than the minimal RXXR motif [19, 22-24]. The released C-terminal fragment is the mature ligand and acts as the signaling molecule [1, 19]. The mature ndr2 ligand has been reported with a range of amino acid sequence lengths, ranging from 119 to 125, due to variability in the cleavage sequence motif and context of amino acid sequence in this region [3, 4, 21].

There are seven cysteine residues that are conserved in most TGF-β family members, six of which interact to form a “cysteine knot” that is essential for biological activity of all TGF-β proteins [25, 26]. The seventh cysteine forms a disulfide bond with a second nodal polypeptide, generating the dimeric form of the ligand that binds to its cognate receptor and leads to activation of downstream signaling pathways [25, 27, 28].

There are 15 previously reported mutations or alleles of zebrafish *ndr2* (https://zfin.org/ZDB-GENE-990415-181) (Supplemental Table 1); eight are chemically-induced mutations and four are radiation-induced mutations. Phenotypic differences in cyclopia severity have been observed across different alleles. Exact sequence information of the mutant alleles is known only for four of these mutant alleles because all of the above alleles were generated and identified using random mutagenesis and phenotypic screening.

This has made any attempt at structure-function studies of proteins arising from all the previously generated mutant alleles difficult. Programmable nucleases have become very useful tools for generating mutations in targeted regions of a gene and the genome, and they enable structure-function studies to be carried out with relative ease. We used the CRISPR-Cas9 nuclease system from *Streptococcus pyogenes* to target distinct regions in all three exons of the zebrafish *ndr2* gene to identify additional alleles.

## Materials and Methods

### CRISPR/sgRNA design and synthesis

CRISPR targets were identified in all three exons of the zebrafish *ndr2* gene (GRCz10, GCA_000002035.3) using a web-based design tool (Benchling [Biology Software], 2017; Retrieved from https://benchling.com.) (Figure 1). Guides were designed and selected across exons with a range of Doench/Benchling scores (range 3–76.82) provided by Benchling (Figure 1; Supplemental Table 2) [29, 30]. Sixteen guide RNA molecules were generated using a cloning-free method previously described [31]. In brief, dsDNA templates with T7 promoter sequences were generated by annealing gene-specific oligos with a constant/tail oligo followed by a fill-in reaction with T4 DNA polymerase (New England Biology, Ipswich, MA) (Supplemental Table 3) [31]. One or two additional G’s were added to the T7 promoter sequence to facilitate efficient transcription (Supplementary Table 3). This product was used as the template for *in vitro* transcription reactions using the T7 Ampliscribe Kit (Epicenter, Madison, WI). Samples were treated with DNaseI and precipitated with ammonium acetate and 100 % ethanol. Air dried samples resuspended in TE buffer were quantified using a Nanodrop UV spectrophotometer and checked for RNA integrity using polyacrylamide gel electrophoresis.

**Figure 1.**
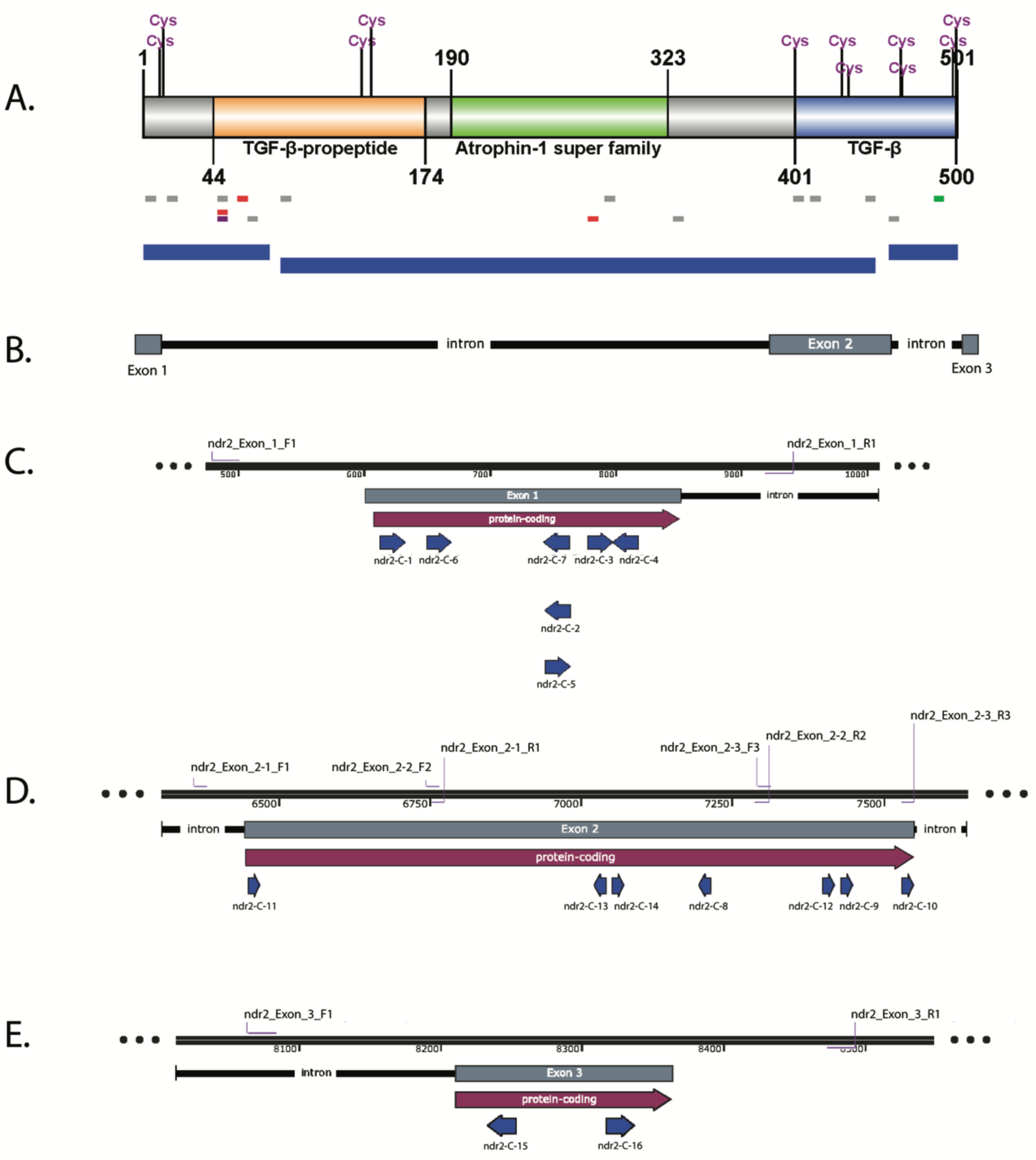
CRISPR targeting in the *ndr2* gene. (A) Schematic showing ndr2 protein structure along with CRISPR sgRNA guides that yielded negative HMA profile and phenotype (gray lines), positive HMA profile and negative phenotype (purple line), negative HMA profile and positive phenotype (green line), positive HMA profile and phenotype (red lines). The bottom blue regions correspond to exons of the *ndr2* gene. (B) Schematic showing *ndr2* gene structure (chr 12: 48,295,205 - 48,302,968). (C-E) Schematics for individual exons (exons 1-3) showing CRISPR targeting regions (blue arrows), PCR primer binding sites (light purple lines), and amplicon sizes.

### In vitro sgRNA-Cas9 nuclease assay

*In vitro* assays for sgRNA-Cas9 nuclease activity were carried out using sgRNAs as previously described [32]. Purified Cas9-NLS protein was obtained from QB3 MacroLab (University of California, Berkeley, CA) and RNP complexes were assembled by combining 10x nuclease buffer (NEBuffer 3.1), Cas9 nuclease (0.1 µL of 2400 ng/uL), sgRNA (0.5 µL of 500 ng/uL) and incubated at 37 °C for 15 minutes. PCR amplicons spanning the CRISPR-targeted regions (Supplemental Table 4) were added to the RNP complex and incubated for 37 °C for one hour followed by RNase treatment at 37 °C for 45 minutes and protein denaturation at 65 °C for 10 minutes. This final reaction was analyzed using gel electrophoresis with 6 % polyacrylamide gels.

### Zebrafish breeding and embryo injections

Zebrafish were housed and maintained at the UAB Zebrafish Research Facility (ZRF). AB strain wild-type zebrafish were used in this study. All experiments were performed in accordance with the recommendations in the *Guide for the Care and Use of Laboratory Animals* published by the National Institutes of Health. The protocols used were approved by and conducted in compliance with the University of Alabama at Birmingham Institutional Animal Care and Use Committee. Cas9 protein–sgRNA RNPs were co-injected into one-cell stage zebrafish embryos. RNP complexes were formed by combining TE lite (10 mM Tris, 0.1 mM EDTA), Cas9 nuclease (0.1 µL of 2400 ng/uL), sgRNA (0.5 µL of 500 ng/uL), and nuclease-free water and incubated at 37 °C for 15 minutes. Each embryo was injected with 3–6 nL of the RNP solution. Dead embryos were removed at mid-gastrulation and the remaining embryos were used for phenotypic analysis and genotyping.

### Phenotypic analysis

Using brightfield light microscopy, 10–20 single injected embryos were examined and embryos showing cyclopia were imaged using a Zeiss Lumar Lamar V12 stereo microscope and AxioVision SE64 Rel.4.9.1 software (Zeiss, Oberkochen, Germany). Embryos were anaesthetized in a solution of embryo medium containing 2 mM tricaine (Sigma-Aldrich, St. Louis, MO) to prevent gross movements during imaging. Embryos were mounted in 3 % methyl cellulose solution (Fisher Scientific, Hampton, NH) to orient them and were imaged in the continued presence of 2 mM tricaine.

### Genotyping using PCR and heteroduplex mobility assay (**HMA**)

Genomic DNA from single embryos was used as the template in polymerase chain reactions (PCRs) with 2x Taq Master Mix (New England Biology, Ipswich, MA). Genomic DNA was obtained by placing single embryos in PCR tubes with 10µL of lysis solution containing Proteinase K and incubated at 55 °C for 2 hours, followed by 95 °C for 10 minutes to inactivate the proteinase K. A small aliquot (0.5 µL) of this solution was directly used as a template to amplify a region flanking each CRISPR target site (Supplemental Table 4). PCR products were analyzed using a heteroduplex mobility assay (HMA) for assessing the nuclease activity and detecting the presence of indels [32-35]. In brief, the amplicons were subjected to denaturation followed by slow renaturation to facilitate the formation of heteroduplexes using a thermocycler. These samples were resolved on 6–8 % polyacrylamide gels and the resulting mobility profiles used to infer the efficiency of CRISPR- Cas9 nuclease activity.

### Targetedamplicon sequence analysis

PCR amplicons obtained from single embryos were cloned into pCR4 using the TOPO-TA cloning kit (Invitrogen, Carlsbad, CA). Ten representative colonies picked from each plate were grown in 1.5 mL liquid cultures to isolate plasmid DNA. Plasmid DNA was sequenced with M13 F and R primers using the Sanger Method.

### Structural modeling

Protein three-dimensional structure predictions of mature wild-type ndr2 (126 amino acids) and translated mutant proteins were made using I-TASSER with no constraints (https://zhanglab.ccmb.med.umich.edu/I-TASSER/) [36-38]; the mature ndr2 amino acid sequence was derived from the precursor protein (nodal-related 2 precursor, 501 amino acids, NCBI Accession: NP_624359.1) following the cleavage recognition sequence of RXRXXR [19, 23] and containing amino acid residues 376–501. Template-based threading and modeling were performed using the top 10 structure templates identified by the Local Meta-Threading-Server (LOMETS) [39], including bone morphogenetic protein-7 (PDB code: 1m4uL), growth factor (PDB code: 5vz3A), TGF-β ligand-receptor complex (PDB code: 3qb4A), and nodal/BMP2 (PDB code: 4n1dA). Images were generated and aligned in the modeling package PyMOL v2.0 (http://www.pymol.org).

## Results

### Invitro assay as a quality control step for nuclease activity

An *in vitro* assay was employed to check for nuclease activity of the sgRNA-Cas9 RNP complexes (Fig. 2A). Guides C-1, C-9, and C-12 did not show any cleavage products suggesting a lack of any nuclease activity. This could be due to degraded or poor sgRNA quality. The extent of nuclease activity for the remaining 12 guides was distinctly visible. The cleavage products correspond to the predicted cut sites in the PCR amplicons. For example, C-2 and C-3 had cut sites in Exon 1 within the same PCR amplicon. Each amplicon had different size cleavage products with C-2 yielding cleavage products of 194 bp and 267 bp and C-3 yielding cleavage products of 146 bp and 315 bp (Fig. 2A). These *in vitro* results showed that 12/15 (80 %) sgRNA guides showed noticeable nuclease activity, providing confidence that they were biochemically active to create double stranded breaks (DSBs) at the target site in the genome of a zebrafish embryo. The GC % of these 12 sgRNA guides that showed clear nuclease activity *in vitro* ranged from 40–85 % (seven were between 40 % and 55 %, six were 60–65 %, one was 70 % and 1 was 85 %). None of the guides had a native 5’ GG start, three had a GN start, and four had an NG start. The length of the guides (21- or 22-mers) and the absence of a native GG sequences did not appear to impact the *in vitro* nuclease activity.

**Figure 2.**
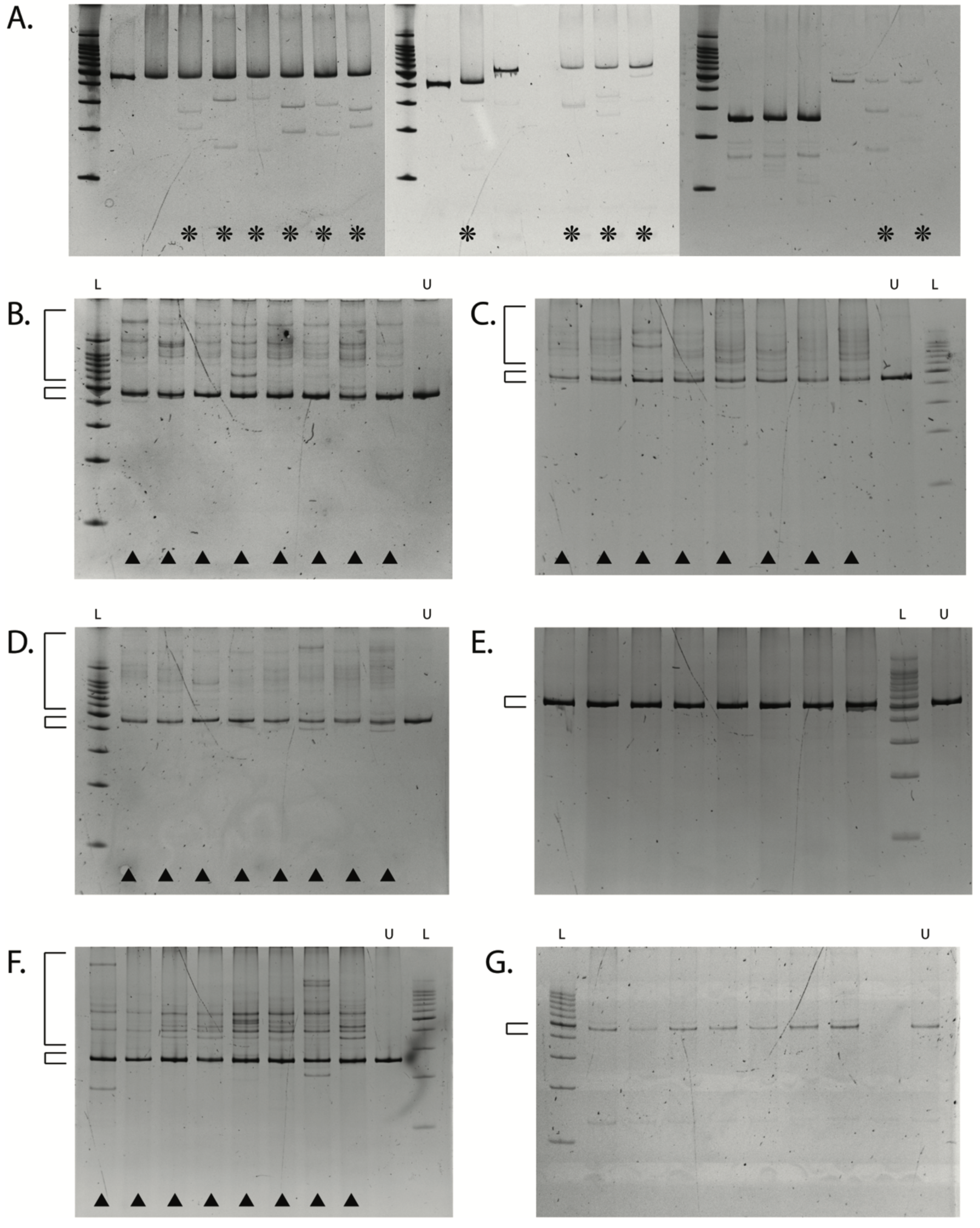
*In vitro* nuclease activity of sgRNA-Cas9 RNP complexes and indel detection in the *ndr2* gene using heteroduplex mobility assay. (A) Images of ethidium bromide stained polyacrylamide gels (6 %) showing *in vitro* sgRNA-Cas9 RNP complex activity by cutting amplified CRISPR-targeted region (positive cutting, star) for all sgRNA. (B-G) Images of ethidium bromide stained polyacrylamide gels (6 %) showing separation of homoduplex and heteroduplex PCR amplicons from sgRNA-Cas9 RNP complexes injected into single zebrafish embryos (positive HMA, arrowheads). CRISPR targeting guide: C-2 (B), C-3 (C), C-7 (D), C-12 (E), C-13 (F), and C-16 (G). Small and large square brackets indicate homoduplex and heteroduplex bands, respectively. L = 100 bp ladder, U = uncut amplified PCR product control or uninjected wild-type control.

### Phenotypic and genotypic analysis of G_0_ embryos injected with sgRNA-Cas9 RNP complexes

Phenotypic analysis showed that 4/12 CRISPR/sgRNAs (33 %) produced penetrant cyclopic phenotypes in 3 days post fertilization (dpf) zebrafish embryos (Fig. 3). Cyclopic mutants exhibited variably fused eyes from mild (Fig. 3B–C) to severe phenotypes (Fig. 3D– E).

**Figure 3.**
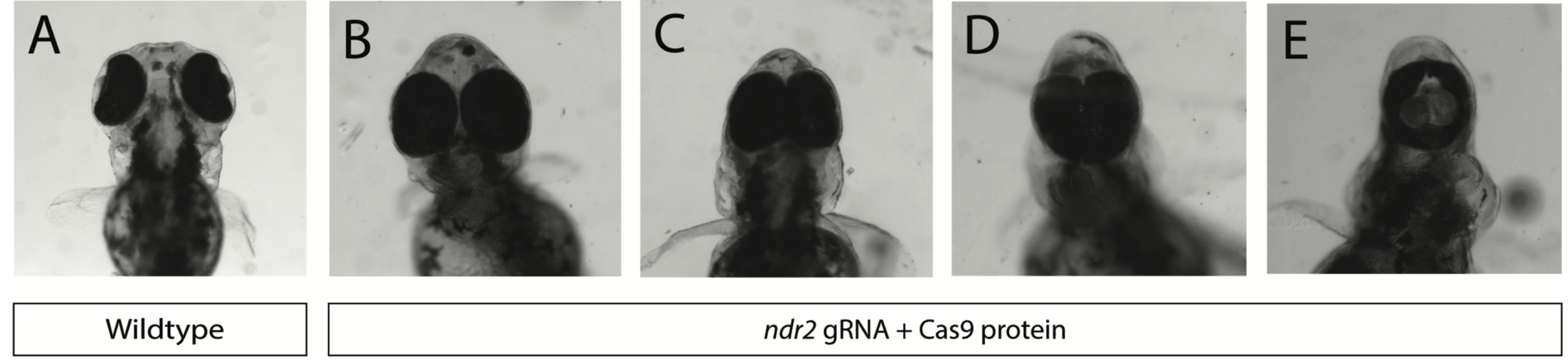
Cyclopia in injected embryos with variable fusion of the eyes. (A-E) Ventral views of wild-type embryo (A) and representative embryos of the mildest phenotypes (B-C) to the most severe phenotypes (D-E) at 3 days post fertilization (dpf).

Genotypic analysis using PCR-HMA indicated that 4/12 CRISPR/sgRNAs (33 %) caused mutations in single embryos that were analyzed (Fig. 2B, C, D, F; Supplemental Table 2). Indels could not be detected using HMA profile with the remaining eight CRISPR/sgRNAs (Fig. 2E & G). There were two instances when the HMA profile did not validate the observed cyclopic phenotype. CRISPR/sgRNA C-2 yielded positive HMA results with no observed cyclopic phenotype (Fig. 2B, Supplemental Table 2), whereas CRISPR/sgRNA C-16 was negative for the HMA profile with a penetrant cyclopic phenotype (Fig. 2G, Supplemental Table 2). An additional round of injections on 32 embryos of the CRISPR/sgRNA C-3 revealed a similar discordant finding between phenotypic and genotypic analyses. Phenotypic analysis showed 2/32 (6 %) injected embryos displaying cyclopic phenotypes. Upon analyzing the same embryos using PCR-HMA, 32/32 (100 %) injected embryos suggested the presence of indels indicated by heteroduplexes (Supplemental Fig 1). Altogether, this suggests that the level of mosaicism and functional outcome of each unique mutation impacts the penetrance and severity of the cyclopic phenotype.

To test the correlation between predicted on-target scores and *in vivo* nuclease activity, we selected sgRNAs with Doench (on-target) scores ranging from 3–77 (Supplemental Table 2). With respect to both the cyclopic phenotype and the nuclease activity detected using PCR-HMA, three of the five active CRISPR/sgRNAs (C-2, C-3, C-7) had Doench (on-target) scores of 63, 62, and 69, respectively. The remaining two active CRISPR/sgRNAs (C-13, C-16) had much lower predictive scores at 19 and 27.9. There were six CRISPR/sgRNAs with predicted scores higher than 30 that did not yield activity (Supplemental Table 2) suggesting that on-target guide scores are not always predictive of *in vivo* nuclease activity.

### Mutations causing loss-of-function phenotypes

Multiple clones from PCR amplicons showing HMA profiles and/or cyclopic phenotypes were Sanger-sequenced to identify the genetic lesions. We obtained 3–6 mutant alleles for the CRISPR target regions (Table 1). Most mutant alleles identified across single embryos from five CRISPR/sgRNAs included small indels, in-frame indels, and complex indels, ranging from 1 to 30 base pairs. There were three larger deletions identified ranging from 71 to 172 base pairs. The majority of the identified mutations, 16/22 (73 %), resulted in frameshifts and predicted premature protein truncation that can explain the loss-of-function phenotypes.

**Table 1.**
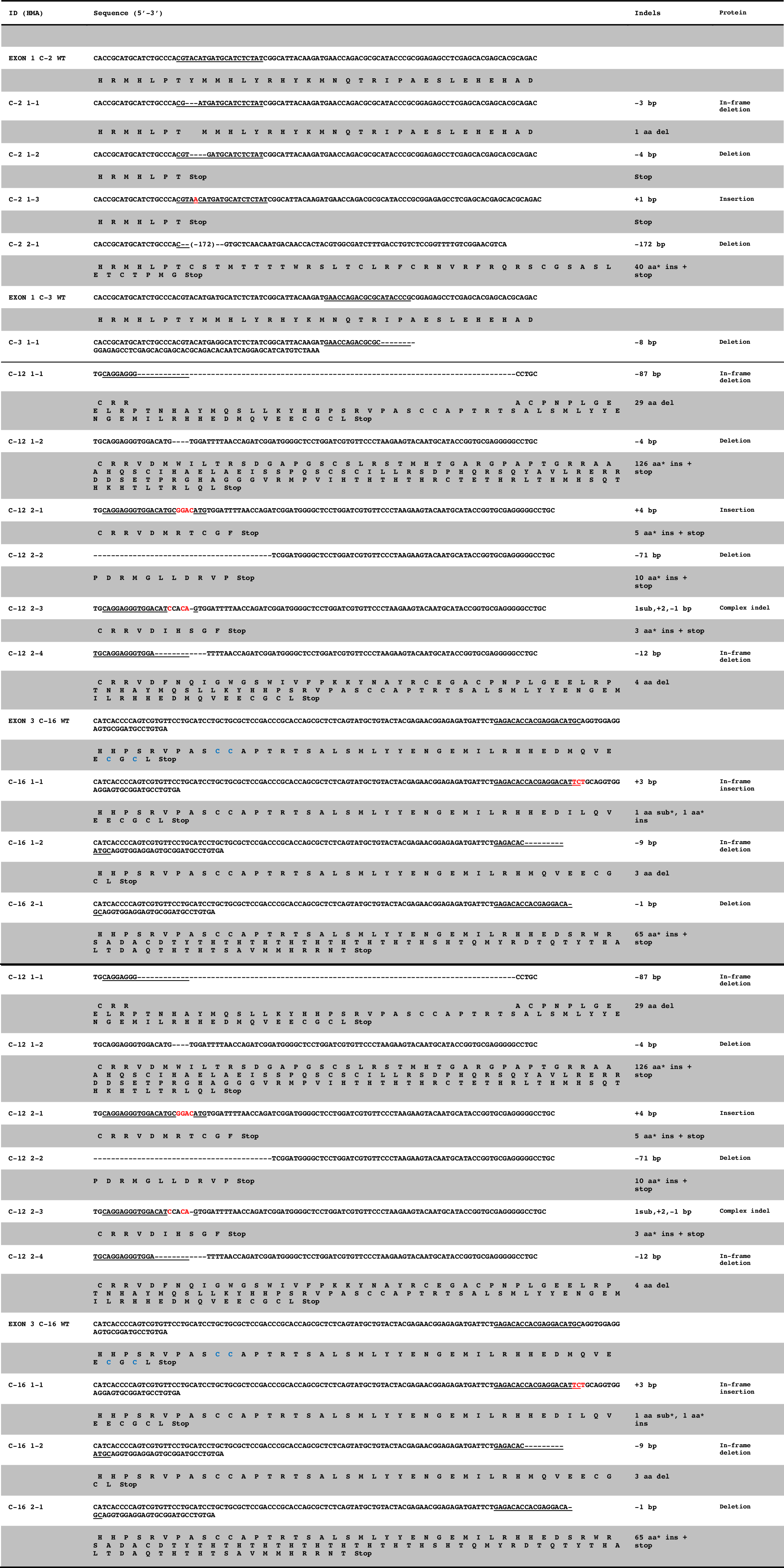
Mutations in *ndr2* induced using sgRNA-Cas9 RNP complexes

As mentioned previously, there were two instances when the HMA profile did not match the observed cyclopic phenotype. For CRISPR/sgRNA C-2 that yielded positive HMA results with no observed cyclopic phenotype, four unique mutant alleles were confirmed with Sanger sequencing. Three of the four mutant alleles caused frameshifts and predicted loss-of-function. With no observable cyclopic phenotype, this suggests that zebrafish embryos injected with C-2 possibly had low contribution of these loss-of-function mutant alleles as compared to the fourth detected in-frame deletion and/or remaining wild-type alleles. For CRISPR/sgRNA C-16 that yielded negative HMA indel profile with a penetrant cyclopic phenotype, three unique mutant alleles were confirmed with Sanger sequencing. One of the three mutant alleles was a 1-bp deletion predicted to cause loss-of-function while the other two yielded in-frame indels. With no observable positive HMA profile and penetrant cyclopic phenotype, this suggests that the zebrafish embryos injected with C-16 possibly had a high contribution of the 1-bp deletion mutant allele that was difficult to resolve using PCR-HMA. These divergent findings from the HMA profile and observed cyclopic phenotypes illustrate the mosaicism of the G_0_ embryos.

### Importance of the “cysteine knot” in ndr2 structure-function

In order to study the structure-function relationship of ndr2 and the guide C-16 mutants, we modeled three-dimensional (3D) structures of a predicted mature ndr2 ligand of 126 amino acids and identified mutant alleles generated from targeting exon 3 (guide C-16) (Fig. 4). I-TASSER provided the following amino acid sequence identities between the mature wild-type ndr2 ligand and corresponding threading templates utilized: 38 % for bone morphogenetic protein-7 (PDB code: 1m4uL), 25 % for growth factor (PDB code: 5vz3A), 42 % for TGF-β ligand-receptor complex (PDB code: 3qb4A), and 62 % for nodal/BMP2 (PDB code: 4n1dA). I-TASSER provided five models for each simulated protein and the top model was selected for further analysis. The I-TASSER confidence score (C-score) for model 1 of the mature wild-type ndr2 ligand was -0.60 with an estimated template modeling score (TM-score) of 0.64±0.13. As observed, the simulated 3D structures of mature wild-type ndr2 exhibited the common elements in the structure of transforming growth factor type β superfamily factors, with five main β-sheets stretching outward structurally from the centrally located “cysteine knot” (Fig. 4A).

**Figure 4.**
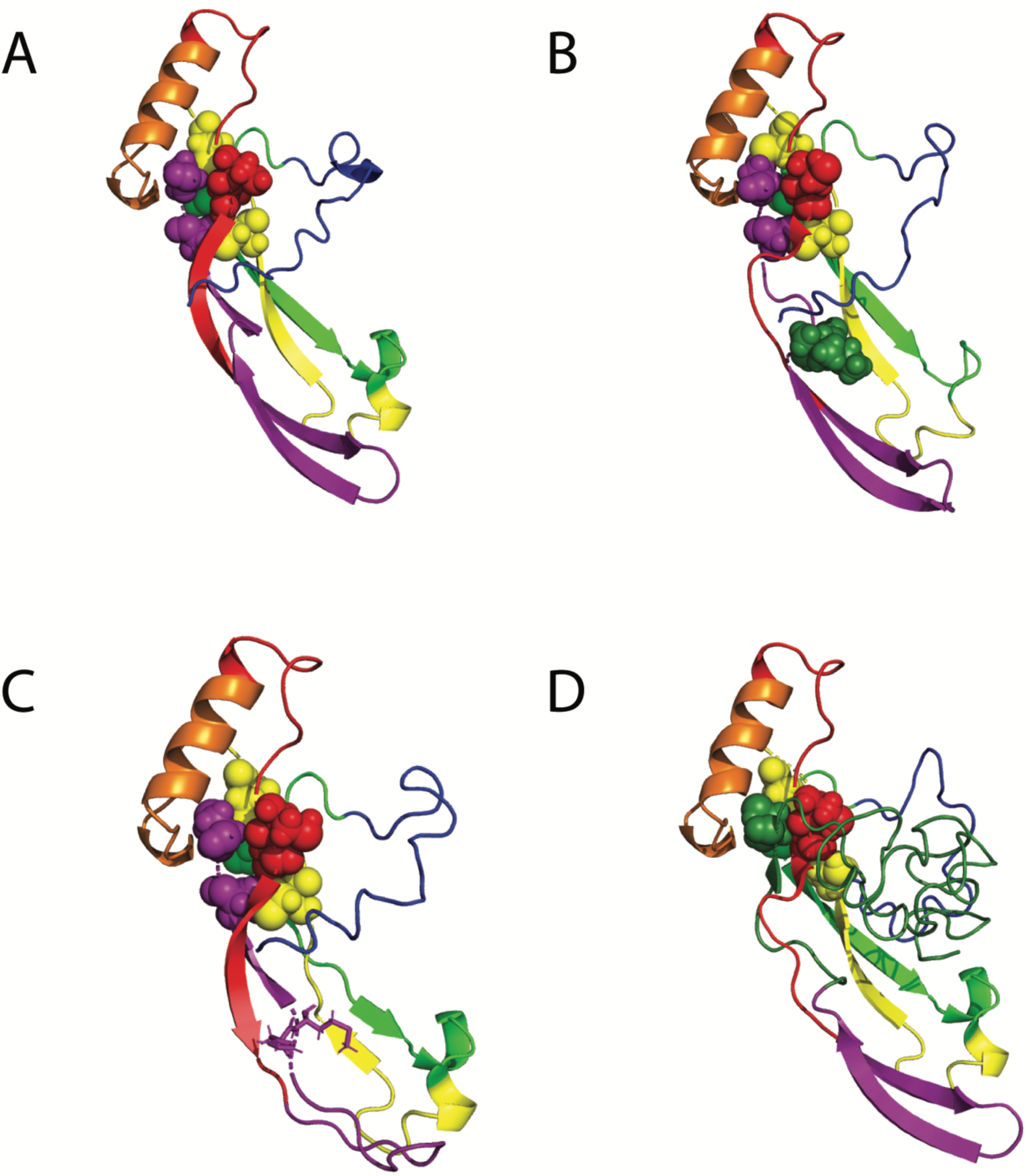
Predicted 3D structure of the mature wild-type ndr2 protein and the mutant alleles generated from targeting exon 3 (guide C-16). (A-D) The structures were generated with ndr2 protein sequence for mature wild type (A), mutant alleles 1-1 and 1-2 resulting in in-frame indels (B-C), and mutant allele 2-1 resulting in a 1 bp deletion (D). Depictions as follows: native cysteine residues are colored spheres (red, yellow, green and purple) (A-D), mutant amino acid residues are dark green spheres (B), two amino acid residues surrounding in-frame deletion are sticks (C), and mutant amino acid residues are dark green ribbon with novel cysteine residue as dark green spheres (D).

The three identified mutant alleles generated from targeting exon 3 (guide C-16) yielded two in-frame indels (models 1-1 and 1-2) and a frameshift deletion (model 2-1). The I-TASSER C-score for the mutant allele 1-1 model was -0.49 with an estimated TM-score of 0.65. The mutant allele 1-1 was identified as a 3 bp in-frame insertion. When we overlaid the protein structures of the mature wild-type ndr2 ligand and the mutant allele 1-1, the overall structure of these two proteins aligned very well with a global root-mean-square deviation (RMSD) of 1.188. The resulting mutant amino acid residue is predicted to disrupt two of the β-sheets resulting in them being smaller than in the wild type and structurally different (Fig. 4B).

The I-TASSER C-score for the mutant allele 1-2 model was -0.65 with an estimated TM-score of 0.63. The mutant allele 1-2 was identified as a 9 bp in-frame deletion. When we overlaid the protein structures of the mature wild-type ndr2 ligand and the mutant allele 1-2, the overall structure of these two proteins aligned very well with a global root-mean-square deviation (RMSD) of 1.383. The loss of three amino acid residues is predicted to disrupt all β-sheets resulting in two different lengths compared to the wild type (Fig. 4C).

The I-TASSER C-score for the mutant allele 2-1 model was -2.65 with an estimated TM-score of 0.41. The mutant allele 2-1 was identified as a 1 bp deletion leading to a frameshift resulting in 65 mutant amino acid residues before terminating the reading frame. When we overlaid the protein structures of the mature wild-type ndr2 ligand and the mutant allele 2-1, structural differences were observed and the RMSD is 1.394. Two native cysteine residues were deleted and one mutant cysteine residue appeared due to this frameshift (Fig. 4D). The mutant cysteine residue was predicted to localize in the “cysteine knot” region and could be functional, but one cysteine residue was still missing. In addition, differences in the β-sheets mentioned above were present. Further, the additional mutant residues formed a long loop that appeared to be highly exposed to solvent. This long loop also sits very close to the cysteine knot.

All identified mutant alleles generated from targeting exon 3 were predicted to cause structural changes to the mature ndr2 ligand that could potentially impact function. Because the position of the native cysteine residues in forming the “cysteine knot” is essential for protein function, we suggest that the mutations we obtained with guide C-16 result were either hypomorphic or null alleles.

## Discussion

Most of the previous mutant alleles of the *ndr2* gene in zebrafish were generated in forward genetic screens with ENU mutagenesis making them difficult to characterize completely at the molecular level (Supplemental Table 1). Using programmable nucleases, especially the CRISPR-Cas9 system, there is a surge of mutant alleles for several genes that can be very well characterized. We tested the efficiency of nuclease activity of CRISPR/sgRNAs with *Streptococcus pyogenes* Cas9 and generated several mutant alleles in the zebrafish *ndr2* gene.

Using a cloning-free method to generate T7-RNA polymerase-driven *in vitro* transcribed sgRNA (modified from [31]) and an *in vitro* nuclease assay, we learned that the lack of native GG dinucleotide at the beginning of the CRISPR guide is not essential for synthesis. As reported earlier, we also observe that guides that are 20–22 nt long can be effective. This is also true with respect to nuclease activity, both *in vitro* and *in vivo*. The *in vitro* nuclease activity is useful as a quality control step prior to embryo injections. However, the presence of *in vitro* nuclease activity did not always translate to *in vivo* activity in embryos. Based on these observations though, we find that the number of usable guides can be more than just the sites that start with canonical GG or GN/NG. In addition, the predicted on-target scores of our sample of CRISPR guides did not always correspond to observed activity in the embryos. This suggests that the current prediction scores for on-target efficiency, especially in the context of zebrafish, need to be further improved using validation experiments. Ultimately, *in vivo* validation of CRISPR guides is necessary.

While Cas9 was delivered as capped mRNA in earlier studies, an increasing number of studies use the Cas9 protein complexed with the guide RNA. Our study validates the effective use of RNP complexes for zebrafish embryo injections, which result in penetrant phenotypes. We were able to observe variable fusion of eyes across injected embryos.

This study also showcases the importance of the “cysteine knot” being essential for biological activity of the ndr2 protein. Even with removal of two native cysteines and the addition of a mutant cysteine in exon 3, loss-of-function and cyclopic phenotypes occurred. Targeting exon 3, we found mutations at the C-terminus of *ndr2* leading to loss of protein function thereby causing cyclopia in zebrafish embryos.

## Data Availability Statement

All relevant data are within the paper and its supporting information files.

## Funding

This work was supported by a Teaching Innovation Grant to AKC by the Quality Enhancement Program (QEP) in the Center for Teaching & Learning, and the Department of Genetics, University of Alabama at Birmingham (UAB). This study was part of a Course-based Undergraduate Research Experience (CURE) for first year undergraduate students in the Science and Technology Honors (STH) Program at UAB. The STH Program provided support for materials and reagents.

## Competing Interests

The authors have declared that no competing interests exist.

## Supplementary Materials

Supplemental Tables (1-4); Supplemental Figure 1

## Acknowledgements

The authors thank the staff of the UAB Zebrafish Research Facility for providing care and maintenance of the facility, Dr. Robert Kesterson and the UAB Transgenic & Genetically Engineered Models (TGEMs) Core facility for support, Dr. Brad Yoder (CDIB, UAB) for providing laboratory space, Dr. Michael Miller (CDIB, UAB) for help with microscopy, Dr. Ben Johnson (VAI) and Dr. David Crossman (Genetics, UAB) for help with sequence analysis, Kartik Manne for advice on structure prediction using PyMol, Ansuya Jogi and Dr. Kiranam Chatti for editing and proofreading, and the administrative support of the Science and Technology Honors (STH) Program. Thanks to Dr. Oreoluwa Adedoyin (MERIT Postdoctoral Scholar) and Lindsay Jenkins (STH undergraduate student) for assistance in the course. AKC thanks Dr. Diane Tucker (STH Program) for advice on designing the CURE.

## Author Contributions

<u>Conceived and designed the experiments</u>: ANT, RSA, IEB, LNB, JCD, DMG, JPG, BAH, ASK, JBM, VLM, ZPM, CLN, RKP, MCR, YAS, APW, SRG, SDF, AKC

<u>Performed the experiments</u>: ANT, RSA, IEB, LNB, JCD, DMG, JPG, BAH, ASK, JBM, VLM, ZPM, CLN, RKP, MCR, YAS, APW, SRG, SDF, AKC

<u>Data analysis</u>: ANT, RSA, IEB, LNB, JCD, DMG, JPG, BAH, ASK, JBM, VLM, ZPM, CLN, RKP, MCR, YAS, APW, SRG, SDF, AKC

<u>Contributed reagents/materials/analysis tools</u>: ANT, RSA, IEB, LNB, JCD, DMG, JPG, BAH, ASK, JBM, VLM, ZPM, CLN, RKP, MCR, YAS, APW, SRG, SDF, AKC

<u>Manuscript preparation</u>: ANT, AKC

